# The spliceosomal component GAMETOPHYTIC FACTOR 1 (GFA1) regulates a key photoperiodic switch

**DOI:** 10.1101/2024.11.01.621476

**Authors:** Sebastian Tiedemann, Lieven Sterck, Hanna Becker, Nicola Nielsen, Cordula Blohm, Daniel Blum, Christa Lanz, Kathrin Maedler, Detlef Weigel, Yves Van de Peer, Rita Groß-Hardt

**Affiliations:** Centre for Biomolecular Interactions, University of Bremen, 28359 Bremen, Germany; Bioinformatics and Systems Biology, VIB/Ghent University, Gent B-9052, Belgium; Centre for Plant Molecular Biology, University of Tübingen, 72076 Tübingen, Germany; School of Science, Constructor University, 28759 Bremen, Germany; Department of Molecular Biology, Max Planck Institute for Developmental Biology, 72076 Tübingen, Germany

## Abstract

Plants have evolved sophisticated mechanisms to perceive and interpret daytime in order to flower at an optimal time point. Here we show that the spliceosomal component *GAMETOPHYTIC FACTOR 1* (*GFA1*) constitutes a previously unrecognized key photoperiodic switch, that is essential for flowering in long days. We show that *gfa1* hypomorphic (*gfa1*_*hyp*_) plants fail to initiate flowering in long-days (LD), which correlates with ectopic activation of the short-day (SD) flowering repressor *ARABIDOPSIS THALIANA CENTRORADIALIS (ATC)*. Accordingly, flowering is restored upon inactivation of *ATC* in *gfa1*_*hyp*_ mutants. A novel tissue-specific *in-planta* splice assay and comprehensive RNAseq profiling of *gfa1*_*hyp*_ mutants indicate that GFA1 mediated pre-mRNA splicing is substrate specific, as previously suggested for GFA1 orthologs. Furthermore, we show that *gfa1*_*hyp*_ mutants accumulate nonsense transcripts of the photoreceptor components *PHYB* and *RRC1*, suggesting inappropriate photoreceptor signaling as a potential cause for the ectopic activation of the SD characteristic profile in *gfa1*_*hyp*_. In fact, known downstream targets of the phytochrome system such as *RS31, SR34a, SRp30* accumulate reduced amounts of light-dependent splice isoforms. Together, our data reveal a link between spliceosome composition and long-day flowering, based on complex transcriptional readouts in response to day length.

## Introduction

Alternative splicing of precursor messenger RNAs (pre-mRNAs) plays a critical role for the development of eukaryotic organisms and greatly enhances protein complexity. In plants, over 40% of all intron-containing genes are alternatively spliced and the majority of the alternative splice isoforms contain premature termination codons (Filichkin *et al*.). It is well established that alternative splicing is regulated by SERINE/ARGININE RICH PROTEINS (SR) and members of the heterogeneous nuclear ribonucleoprotein (hnRNP) family (Nilsen *et al*.). It was only in 2007 that results in budding yeast, *Saccharomyces cerevisiae*, have shown that spliceosomal components are unequally involved in the removal of non-coding sequences. Since then, defects in various core spliceosomal components have been shown to differentially affect transcriptional readout.

In the plant *Arabidopsis thaliana*, alternative splicing and the function of different core spliceosomal proteins have for example been demonstrated to be important for female gametophyte development and flower organ growth (Groß-Hardt *et al*., Yagi *et al*.). We have previously shown that loss-of-function mutations in the genes for the core spliceosomal proteins LACHESIS (LIS), GAMETOPHYTIC FACTOR 1 (GFA1) and ATROPOS (ATO) cause the formation of supernumerary egg cells, suggesting that spliceosome-regulated pre-mRNA splicing is critical for the regulation of gametic cell fate (Groß-Hardt *et al*., Moll *et al*.). *GFA1* is also involved in embryogenesis and contributes to the directional growth of floral organs (Liu M. *et al*., Yagi *et al*.). *GFA1* encodes the *A. thaliana* ortholog of the *S. cerevisiae* pre-mRNA splicing factor *Snu114*. Snu114 is associated with the U5 subunit and is required for the activation and degradation of the spliceosome (reviewed in Frazer *et al*., Jia *et al*.) Disruptions of orthologs in human, mouse, zebrafish, and frog lead to severe developmental defects and mis-splicing events (Beauchamp *et al*., Lei *et al*., Park *et al*.). Previous studies in plants have suggested a conserved role of *GFA1* in pre-mRNA splicing; GFA1 interacts with the putative U5 small nuclear ribonucleoprotein (snRNP) components BRR2a and PRP8, co-localizes with the serine/arginine rich pre-mRNA splicing factor SC35 (Liu M. *et al*., Yagi *et al*.) and localizes to nuclear speckles (Moll *et al*.). Here we show that reduced GFA1 activity differentially affects intron removal within and between individual transcripts, suggesting that GFA1 regulates pre-mRNA splicing in a substrate specific manner. In addition, we show that GFA1 is necessary for the integrity of the phytochrome system and the SD-to-LD switch, and that this step is regulated by GFA1-dependent repression of *ARABIDOPSIS THALIANA CENTRORADIALIS (ATC)*.

## Results and Discussion

### A novel plant germline splice assay suggests substrate specificity for core spliceosomal component GFA1

The *gfa1* mutant was initially identified on the basis of ectopic egg cell marker expression (Moll *et al*.), but the late and specific arrest shortly before fertilization is difficult to reconcile with a general role of GFA1 in pre-mRNA splicing.

To determine whether GFA1 is required for every pre-mRNA splicing event, we in a first step developed an *in-planta* splice assay, which allows monitoring of intron removal in *gfa1* female gametophytes. For this purpose, we generated minigenes by amplifying exon-intron-exon cassettes of selected genes. The minigenes were introduced in frame between an N-terminal nuclear localization signal and a C-terminal *GFP* reporter (*NLS_e-i-e_GFP*). We chose the fragments such that intron retention resulted in a frame shift, preliminary stop codons, or both, with the rationale that GFP is only detected upon intron removal. The *GFP* minigenes were expressed from the *pAt5g40260* promoter, which confers expression in the female gametophyte but not in the surrounding sporophytic tissue (Yu H.J. *et al*). In wild-type female gametophytes, *pAt5g40260::NLS_GFP* plants exhibit a prominent GFP signal in the central cell and weak expression in other female gametophytic cells (Fig. 1A). Synergids and central cells in *gfa1* female gametophytes become reprogrammed into cells expressing a molecular profile characteristic to egg cells, whereas antipodal cells express central cell characteristic features (Groß-Hardt *et al*., Moll *et al*.). This reprogramming is accompanied by changes in nuclear position and a failure of the central cell nuclei to fuse (Moll *et al*.). As a consequence, the intron-free *pAt5g40260::NLS_GFP* reporter is nearly uniformly detected in all cells of *gfa1* female gametophytes (Fig. 1B).

**Figure 1:**
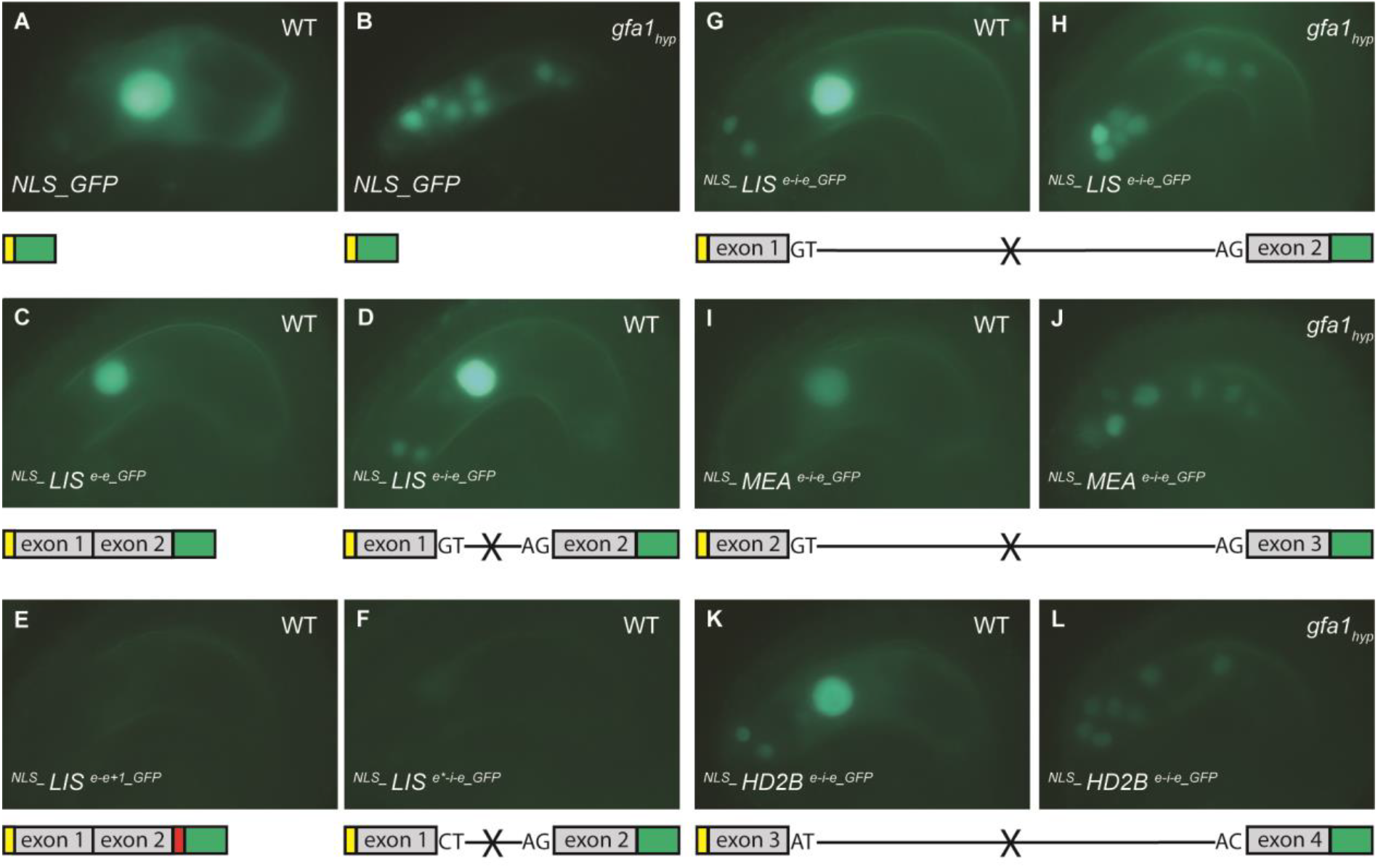
*In-planta* splice assay probing canonical and non-canonical splice sites in wild-type and *gfa1* female gametophytes. (A-F) Control constructs: (A) *NLS_GFP* in wild-type and (B) *gfa1* mutants, (C) two consecutive exons of *LIS*, (D) same exons as (C) interrupted by an intron. (E) Same construct as (C) with an additional frame shift inducing base pair 3’ of the second exon. (F) Same construct as (D) with mutated 5’ splice site to suppress intron removal. (G-L) Different GFP-minigenes in wild-type (G, I, K) and *gfa1* female gametophytes (H, J, L,). (G, H) *NLS_ LIS e-i-e_GFP*, (I, J) *NLS_ MEA e-i-e_GFP*, (K, L) *NLS_ HD2B e-i-e_GFP*, containing a non-canonical splice site. construct structure is depicted below and not to scale. Exons, grey bar; introns, black lines; *NLS*, yellow bar; *GFP*, green bar; frame shift, red bar. Presence of multiple stop codons that would result from intron retention are indicated by X. Female gametophytes were characterized 2.5 d after emasculation.

To test the validity of the *in-planta* splice assay, we first introduced various control constructs into wild-type plants and characterized their expression: As expected, expression of an *NLS_ LIS e-i-e_GFP* construct in wild type yielded comparable results to an intron-free construct (Fig. 1C, D). Importantly, the GFP signal was abolished when a frame shift mutation 5’ of the *GFP* was introduced (Fig. 1E). Likewise, no GFP was detected following disruption of the 5’ splice site through site-directed mutagenesis (Fig. 1F), indicating that the *in-planta* splice assay reliably detects intron retention. We next introduced three minigenes probing recognition of consensus and non-consensus splice sites into *gfa1/GFA1* plants. The *GFP*-minigenes yielded different signal intensities in wild-type female gametophytes (Fig. 1G, I, K,), which might reflect differences in protein stability or a requirement for genomic splicing enhancers/silencers not included in the construct. Surprisingly, in *gfa1* mutant female gametophytes containing *NLS_LIS_e-i-e_GFP* and *NLS_MEDEA(MEA)_e-i-e_GFP*, we observed substantial GFP signals indicative of correct intron removal at a time point when morphological defects were clearly evident (Fig. 1G-J, compare Fig. 1A, B). Similarly, a splicing cassette for *HISTONE DEACETYLASE 2B* (*NLS_HD2B_e-i-e_GFP*), which contains a non-canonical AT-AC splice site, was correctly processed (Fig. 1L). Given the restricted sensitivity of the splicing assay we do not rule out that splicing of the minigenes is impaired to some extent. However, our results are not compatible with a universal role of GFA1 in pre-mRNA splicing but rather suggest substrate specificity. In order to better understand the mode of GFA1-directed mRNA processing, we analyzed GFA1 dependent pre-mRNA splicing in sporophytic tissue.

### *gfa1* hypomorphic mutants fail to initiate flowering under LD conditions

While *gfa1* heterozygous plants resemble wild type in their growth and morphology, homozygous *gfa1* mutant plants cannot be recovered (Moll *et. al*). We therefore decided to generate *gfa1* hypomorphic *(gfa1*_*hyp*_) plants by expressing the cDNA of *GFA1* under the weak promoter of the *GFA1* paralog *GAMETOPHYTIC FACTOR 1 LIKE (GFL)* in *gfa1* mutants (Coury *et al*.). While the expression of GFA1 under its endogeneous promoter expression rescues the mutant (Moll *et. al*), *gfa1*_*hyp*_ plants showed several defects already in the vegetative phase, including being small, with serrated and curled leaves (Fig. 2A) and having a substantially shortened root (Fig. S1). In addition, these plants show daylength-dependent flowering defects, as described below. A control expressing the partial complementation construct in wild type (*pGFL::cGFA1/*+) did not show any obvious abnormalities (Fig. 2A), indicating that the phenotypes are not due to a dominant effect of the transgene, but most likely due to reduced GFA1 activity because of a reduction in *GFA1* transcripts. Expression of an additional *GFA1* copy in wild type (*pGFA1::cGFA1/*+) did not cause any obvious defects, supporting the assertion that the phenotypes of the *gfa1*_*hyp*_ plants are not due to dominant gain-of-function effects (Fig. 2A).

**Figure 2:**
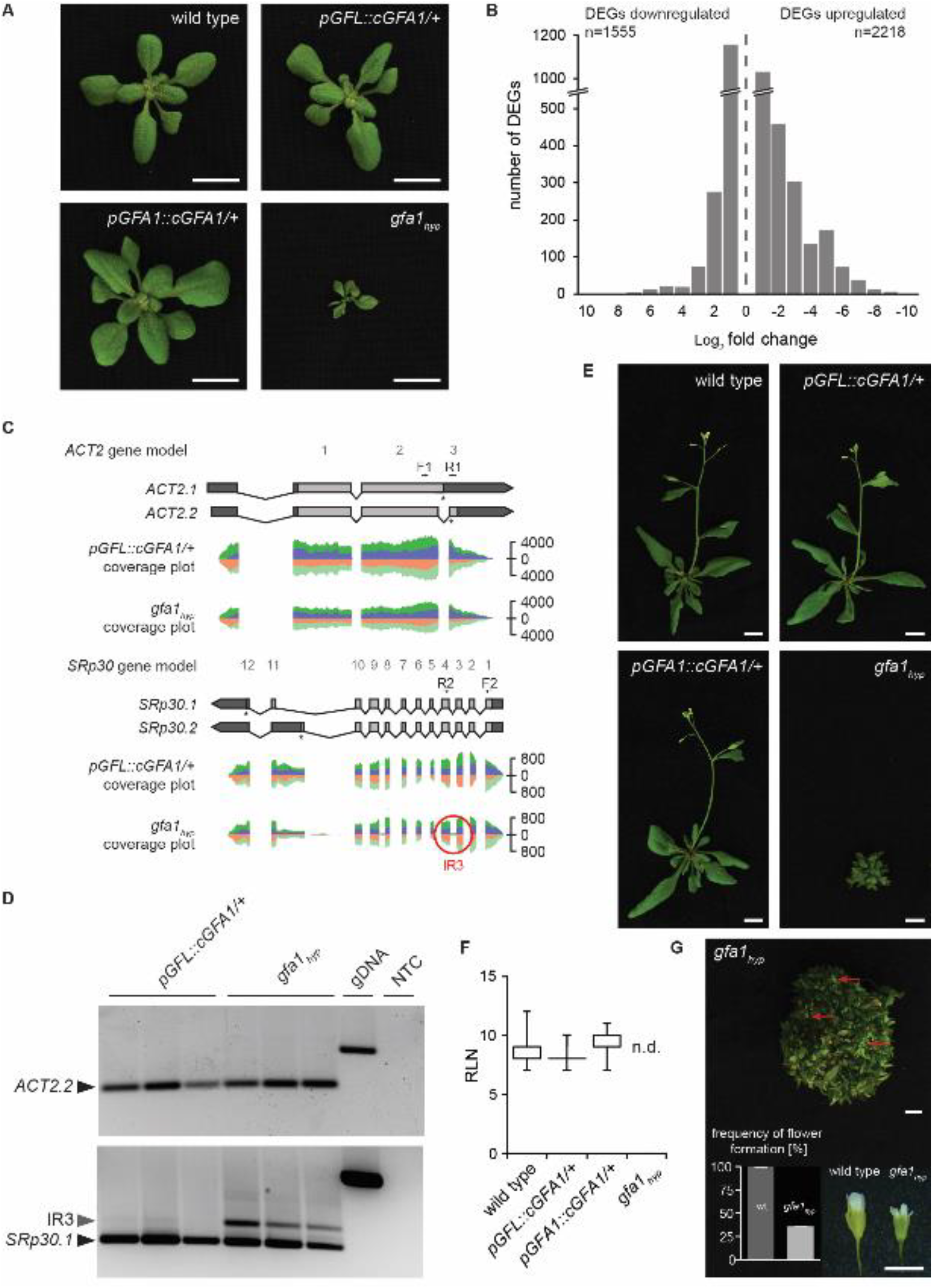
RNA-seq analysis of GFA1 function in *gfa1*_*hyp*_ plants. (A) 19-day old *gfa1*_*hyp*_ plants in comparison to wild-type, *pGFL::cGFA1/*+, and *pGFA1::cGFA1/*+ plants. Scale bar: 1 cm. (B) Significantly differentially expressed genes (DEGs) in 12-day old *gfa1*_*hyp*_ seedlings in comparison to *pGFL::cGFA1/*+ seedlings (FDR<0.05 and log_2_FoldChange>1), ranked by fold change. (C) TAIR10 gene model and RNA-seq coverage in 12-day old *pGFL::cGFA1/*+ and *gfa1*_*hyp*_ seedlings for *ACT2* and *SRp30*. Introns represented by black lines and exons by light grey boxes. UTRs are represented by dark grey boxes. Stop-codon labeled by asterisks. Reads are colored by forward (dark green/orange) and reverse strand (purple/light green) for sense and antisense transcripts, respectively. IR3 is highlighted with a red circle. Arrowheads indicate positions of forward (F) and reverse primers (R) for RT-PCR analysis in (D). (D) RT-PCR analysis of *ACT2*.*2* and *SRp30*.*1* transcripts in 12-day old *pGFL::cGFA1/*+ and *gfa1*_*hyp*_. Genomic DNA (gDNA) was considered as positive, no-template control (NTC) with distilled water as negative control. (E) 23-day old wild-type, *pGFL::cGFA1/*+, *pGFA1::cGFA1/*+, and 40-day old *gfa1*_*hyp*_ plants at 23°C LD. Scale bar: 1 cm. (F) Flowering time scored as rosette leaf number at 0.5 cm shoot elongation. n.d., plants did not show shoot elongation by the end of the experiment, 6 weeks after germination (see also Table S2A). (G) Some *gfa1*_*hyp*_ plants flower without shoot elongation 97 days after germination. Red arrows point to flowers. Frequency of flower formation of wild-type (n=76; dark grey) and *gfa1*_*hyp*_ (n=84; light grey) plants 65 days after germination determined for three independent experimental replicates in 23°C LD (see also Table S2 A, B, C). Scale bar: 0.5 cm. Error bars, SEM. For all plant pictures brightness was equally increased in Adobe Photoshop.

### mRNA profiling of *gfa1* hypomorphic mutants confirms substrate specific role of GFA1 in pre-mRNA splicing

To get an idea on how extensive the substrate specific effects of GFA1 are, we first determined the transcriptional landscape of 12-day old *gfa1*_*hyp*_ seedlings by RNA-seq. Transcriptional profiling based on the transcript isoform of the main splicing isoform annotated in TAIR10 corresponding to exonic regions and splice junctions conducted by the 5’ transcript start and 3’ transcript end of a gene. In *gfa1*_*hyp*_ plants, the amount of *GFA1* transcripts was only half of that in *pGFL::cGFA1/*+ control plants (Table S1B, Fig. S2A, C), with the *GFA1* paralog *GFL* being upregulated (Table S1C), indicating that *GFL* is under regulatory control by GFA1 (Fig. S2B, C).

When analyzing the main splicing isoforms of all expressed genes, we identified among the over 20,000 detectable genes (Table S1A) nearly 4,000 differentially expressed genes (FDR<0.05 and log_2_FoldChange>1), with 1,555 genes being down-and 2,218 genes being upregulated (Fig. 2B and Table S1B, C). The majority of differentially expressed genes presented log_2_FoldChanges between ±1 and ±2 (Fig. 2B). Only seven genes reached a log_2_FoldChange over ±9 (Fig. 2B). In support of the *in-planta* splice assay, we identified correctly processed transcripts such as *ACT2*.*2*, which is the predominant splice isoform of *ACT2* (Fig. 2C).

To identify mis-spliced genes, we searched for upregulated pre-mRNA reads in the *gfa1*_*hyp*_ transcriptome that were aligned to intronic regions (FDR<0.05 and log_2_FoldChange>1, Table S1D). We focused on transcripts with increased intron reads in *gfa1*_*hyp*_ plants compared to the control plants. We identified intron retention events (IR) in genes with a single intronic region, such as *EIN3-BINDING F BOX PROTEIN 1 (EBF1*.*1)*, and in genes with multiple introns, such as *1-AMINOCYCLOPROPANE-1-CARBOXYLIC ACID SYNTHASE 6 (ACS6*.*1)* (Table S1D and Fig. S3A). In the latter category, we also found genes where splicing was only affected in a subset of introns (Fig. S3B). Transcripts, such as *MEDIATOR OF RNA POLYMERASE II TRANSCRIPTION SUBUNIT 19A-LIKE PROTEIN (MED19A*.*1)*, had IR events only in their 5’ UTR or 3’UTR region as well as both UTR’s, whereas all introns within the coding region of the gene were spliced correctly (Fig. S3B). *CONSTITUTIVE TRIPLE RESPONSE 1 (CTR1)* and *EARLY RESPONSE TO DEHYDRATION 6 (ERD6)* transcripts contained a large number of intronic regions within the coding sequence. However, only a few introns were not properly removed by splicing (Table S1D and Fig. S3B). These data indicate that GFA1 is not equally required for all pre-mRNA splicing events. We were specifically interested in IR events in mRNAs coding for known pre-mRNA splice regulators. While they did not necessarily fulfill our significance criteria, IR hinting reads were detected e.g. for *SERINE/ARGININE-RICH RPOTEIN SPLICING FACTOR 30 (SRp30)* (Fig. 2C). Hence, we confirmed this IR event by RT-PCR using specific intron-spanning primers (Fig. 2D). IR events were clearly detectable in *gfa1*_*hyp*_ but not control plants (Fig. 2D). This approach also confirmed correct splicing of the *ACT2*.*2* transcript (Fig. 2D). Similar results were obtained for *ARGININE/SERINE-RICH SPLICING FACTOR 31* (*RS31*) and *SC35-LIKE SPLICING FACTOR 33* (*SR33*) (Fig. S4). Noteworthy, all validated IR events result in transcripts with premature termination codons.

We have only scratched the surface when it comes to our understanding of the differential regulation of pre-mRNA splicing by core-spliceosomal components. Here, we show that reduction of the spliceosomal component GFA1 affects transcript processing. Importantly, the splice modifications are not a simple function of expression level in *gfa1*_*hyp*_ mutants, with IR often affecting only individual introns in a transcript. In agreement with previous work in yeast (Pleiss *et al*.), our results suggest that the spliceosome is a modular molecular machine and that spliceosomal composition critically determines the composition of the transcriptome.

### GFA1 is necessary for correct processing of photoreception associated genes

RNA-seq data showed that several introns were improperly spliced in PHYB and RRC1 (Table S1D). RT-PCR confirmed enhanced IR2 and IR3 events in RRC1 (Fig. 3A, B), and IR3 and IR2 events in PHYB in *gfa1*_*hyp*_ plants, both in 23°C and 27°C LD (Fig. 3C, D). All resulting IR transcripts contain stop codons. PHYA/PHYB and RRC1 regulate alternative splicing in a light dependent manner (Shikata1 et al., Shikata2 et al., Hartmann et al.). Some targets of this regulation, i.e. the splicing factors RS31, SR34a, and SRp30, show an altered splice isoform profile upon *GFA1* reduction at 23°C and 27°C (Fig. 3E).

**Figure 3:**
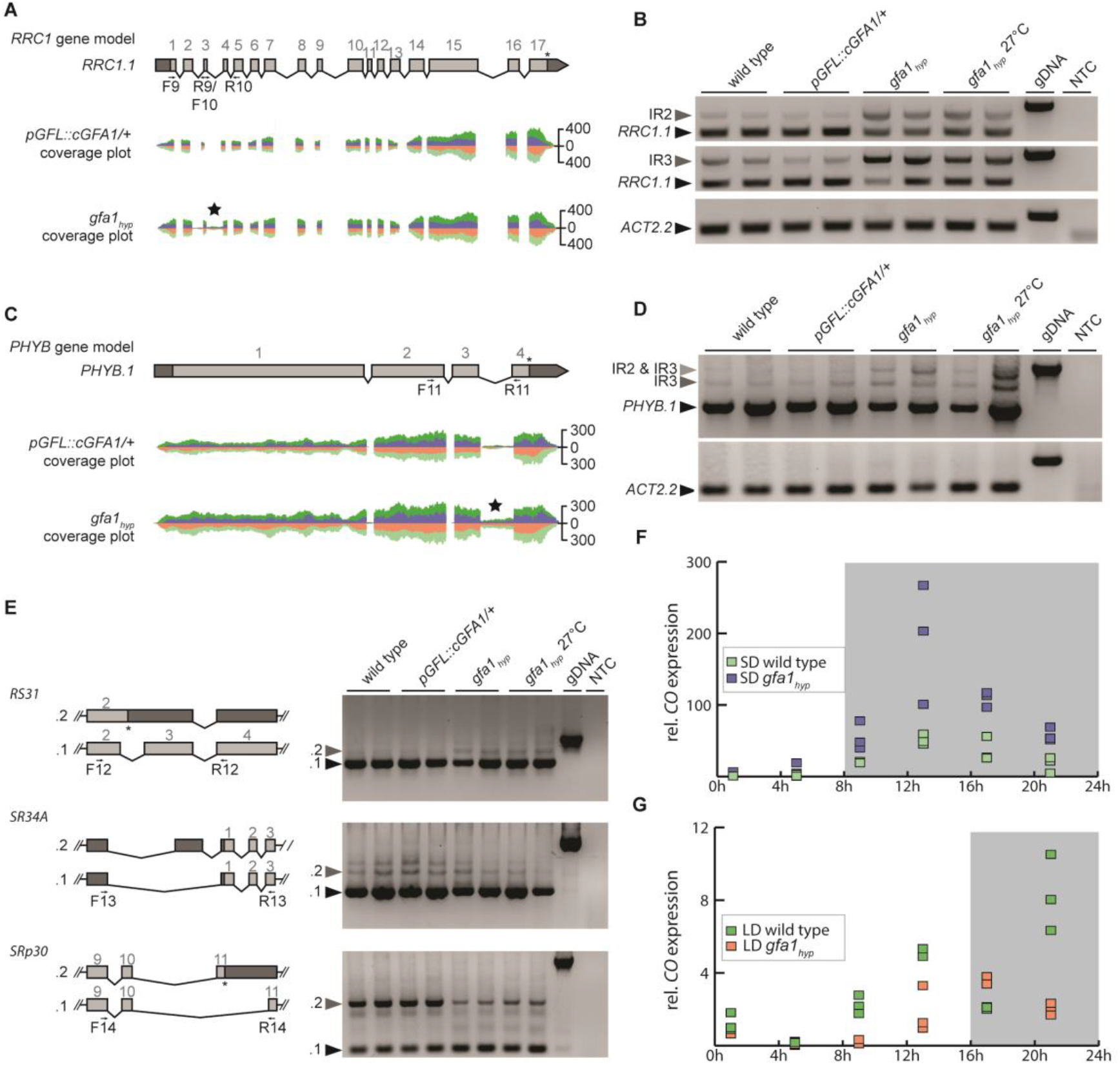
Enhanced intron retention in *RRC1, PHYB* and *SR* genes and LD specific changes in *CO* expression. TAIR10 gene model and RNA-seq coverage in 12-day old *pGFL::cGFA1/*+ and *gfa1*_*hyp*_ seedlings for *RRC1* (A) and *PHYB* (C). Black asterisk indicates significantly increased intron coverage in *gfa1*_*hyp*_ in comparison to *pGFL::cGFA1/*+ seedlings (FDR<0.05 and log_2_FoldChange>1). RT-PCR on *RRC1* IR2 and IR3 (B) and *PHYB* IR3 (D) event in 20-day old plants. *ACT2*.*2* serves as loading control. Arrowheads indicate positions of forward (F) and reverse primers (R). Gene model and RT-PCR analysis of light-dependent alternative splicing variants of different *SR* genes. Introns are represented by black lines and exons by light grey boxes. UTRs are represented by dark grey boxes. Premature stop-codon labeled by asterisks. Arrowheads indicate positions of forward (F) and reverse primers (R). Genomic DNA (gDNA) was used as positive, no-template control (NTC) with distilled water as negative control. (F and G) Relative *CONSTANS* expression over 24h in non-bolting wildtype and *gfa1*_*hyp*_ plants grown under SD (F) and LD (G) as determined by qPCR, grey areas indicate dark phases; *ACT2* and *UBQ10* are used as a reference and the expression is presented relative to one data point at 1h.

The splicing defects in photoreception components *PHYB, RRC1* and downstream factors, prompted us to investigate if photoregulation is deviating in *gfa1*_*hyp*_ plants. Photoreception is key for the alignment of the circadian clock. Hence, we examined mRNA levels of two central circadian clock oscillators, *CCA1* and *TOC1*, over 24h. We observed no shift in the overall pattern between *gfa1*_*hyp*_ and WT plants (Fig. S12 C-F). Furthermore, *GFA1* mRNA level appears to not be subjected to circadian fluctuations (Fig. S12A, B). A key physiological function of photoreception is conditional induction of flowering. By assessing *CONSTANS* (*CO*) mRNA levels over 24h as an indicator for photoinduction of flowering, we found differences between WT and *gfa1*_*hyp*_ plants. Under long day conditions (LD), WT plants exhibit two peaks around dusk (Fig. 3G). This pattern was not detected in *gfa1*_*hyp*_ plants, which instead exhibit a single peak around dusk under LD conditions (Fig. 3G). Comparable to WT, a single expression peak is found in *gfa1*_*hyp*_ plants under short day (SD) conditions after dusk (Fig. 3F). The specific molecular defect observed under LD conditions indicates that GFA1 functions in the integration of LD stimuli and concomitant activation of flowering.

### GFA1 enables flowering under LD conditions through downregulation of the dominant flowering repressor ATC

The altered expression pattern of the flowering inducer *CO* specifically under LD conditions prompted us to investigate flowering under LD and SD conditions in *gfa1*_*hyp*_ : While wild-type plants and plants expressing *pGFL::cGFA1/*+ or *pGFA1::cGFA1/*+ flowered in LD with about nine rosette leaves (Fig. 2E, F), *gfa1*_*hyp*_ plants fail to induce flowering before the experiment was terminated after 6 weeks (Fig. 2E, F); a similar behavior was observed under continuous light (Fig. S5). After 9 weeks, only 33% of *gfa1*_*hyp*_ plants eventually formed flowers (Fig. 2G), but the main shoot never elongated, even when *gfa1*_*hyp*_ plants were kept for 14 weeks (Fig. 2G). However, flower formation occurred significantly delayed in days after sowing in comparison to the controls (Fig. S5B). Due to strong g*fa1*_*hyp*_ leaf phenotype, we were not able to clearly discriminate between rosette and cauline leaves. Nevertheless, 45-day old *gfa1*_*hyp*_ plants formed more than 200 leaves without showing visible flowers, which is at least 5-times more than the total leaf number generated by wild-type, *pGFL::cGFA1/*+ *and pGFA1::cGFA1/*+ plants at 0.5 cm shoot elongation (Fig. S5C).

*gfa1*_*hyp*_ thus is unique among late flowering mutants, where failure to flower at all is rare, except in plants with very high levels of expression of the flowering repressor *FLC* (Sheldon *et al*., Michaels *et al*.). However, failure to flower in *gfa1*_*hyp*_ is not due to *FLC* overexpression. On the contrary, *FLC* expression is greatly reduced (Fig. S6A and Table S1B). Expression of other genes of the vernalization and autonomous pathway, in which *FLC* plays a central role (reviewed in Bloomer and Dean *et al*.), was also unaffected (Fig. S6A and Table S1A). In agreement, vernalization did not accelerate flowering of *gfa1*_*hyp*_ mutants (Fig. S6B).

Apart from vernalization, elevated ambient temperature can greatly accelerate flowering of many *A. thaliana* genotypes (Balasubramanian *et al*.), and this was also observed for *gfa1*_hyp_ plants, which started to flower with an average of 9 rosette leaves at 27°C LD, which was only slightly later than the control plants (Fig. S7A, B). Given that GFA1 reduction inflicts wide-ranging splicing defects, a prime candidate for a gene releasing the *GFA1* flowering block in a temperature-dependent manner is the flowering repressor *FLOWERING LOCUS M* (*FLM)*. Temperature changes the ratio between two main splice isoforms, the flowering repressor *FLM-β* and the flowering activator *FLM-δ* (Scortecci *et al*., Balasubramanian *et al*., Scortecci *et al*., Posé *et al*.). To test whether a change in *FLM* splice isoforms can indeed bypass the *gfa1*_*hyp*_ flowering defects, we introduced an *flm-3* mutant allele into *gfa1*_*hyp*_ and separately overexpressed *FLM-δ* in *gfa1*_*hyp*_. While the *gfa1*_*hyp*_ leaf phenotype was unaffected (Fig. S7C), both approaches induced flowering in *gfa1*_*hyp*_ in 23°C LD, with *gfa1*_*hyp*_ *flm-3* plants flowering as early as wild-type plants (Fig. S7D, E). Investigating if *FLM* splicing may be causal for the late flowering of *gfa1*_*hyp*_ mutants, we compared *FLM-β* and *FLM-δ* transcript abundance in 20-day-old plants at 23°C and 27°C by qPCR. We did not observe substantial changes in *gfa1*_*hyp*_ (Fig. S8B), indicating that the *gfa1*_*hyp*_ defects are independent of *FLM*. Altogether, these observations suggest that *GFA1* functions in a temperature-independent pathway of flowering repression.

That *GFA1* affects a separate flowering regulatory pathway is supported by RNA-seq data, which did not point to major changes in the expression of most key flowering regulators (reviewed by Johanssen and Staiger *et al*.) of the gibberellin, autonomous and photoperiodic pathways (Fig. S9). A striking exception was *ATC* (Huang *et al*.) (Fig. 4A, B; Fig. S9). ATC is a potent SD flowering repressor and its overexpression in LD causes plants to flower late, presumably by interfering with formation of the FLOWERING LOCUS T/FLOWERING LOCUS D (FT/FD) flowering activator complex (Huang *et al*.). *CO, FT* and *FD* were overall expressed in *gfa1*_*hyp*_ plants (Fig. S10). *GFA1* expression as determined by qPCR was not significantly altered between LD and SD (foldchange below 1). Consistent with *ATC* overexpression being responsible for late flowering of *gfa1*_*hyp*_ plants, the *atc-2* knock-down allele greatly accelerated flowering of *gfa1*_*hyp*_ plants (Huang *et al*.) (Fig. 4), without greatly affecting leaf or root morphology of *gfa1*_*hyp*_ plants (Fig. 4D; Fig. S11). Since GFA1 is necessary to release repression of flowering by *ATC* in LD, we wanted to learn whether *GFA1* constitutes a genuine day length switch. We found that plants containing an extra *GFA1* copy flower early in SD, indicating that *GFA1* can positively and negatively influence flowering time (Fig. 4F, G).

**Figure 4:**
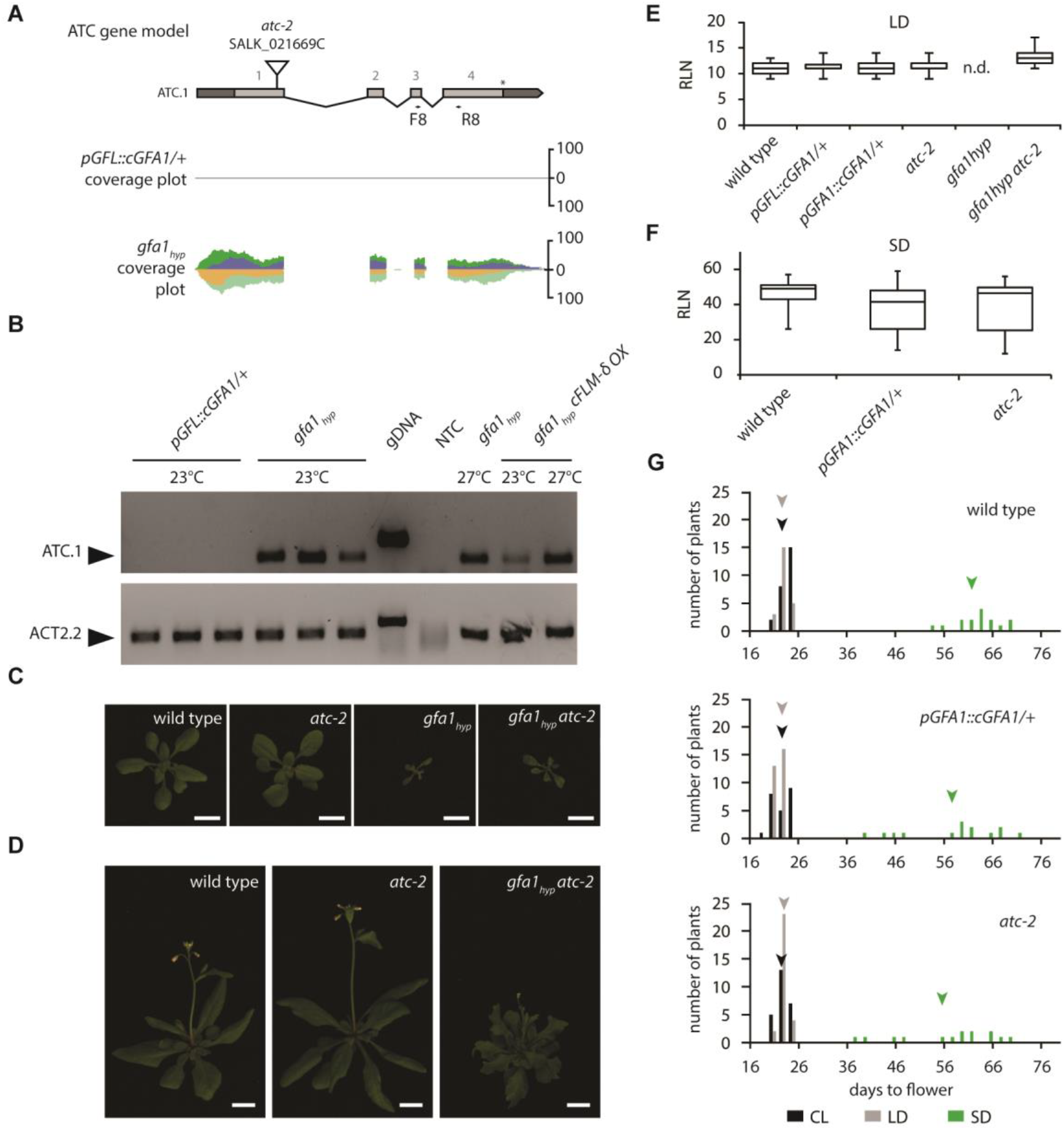
*ATC* overexpression delays flowering of *gfa1*_*hyp*_ and extra *GFA1* accelerates flowering under SD (A) *ATC* TAIR10 gene model and RNA-seq coverage in 12-day old *pGFL::cGFA1/*+ and *gfa1*_*hyp*_ seedlings. The position of the T-DNA insertion in the knock-down allele *atc-2* is shown above the gene model. For *pGFL::cGFA1/*+, the number of reads was too low to be depicted. (B) RT-PCR analysis of *ATC* expression in 20-day old *pGFL::cGFA1/*+, *gfa1*_*hyp*_ and *gfa1*_*hyp*_ *cFLM-δ* OX plants grown in LD at 23°C or 27°C. *ACT2*.*2* transcript serves as loading control. Arrowheads indicate positions of forward (F) and reverse primers (R). Genomic DNA (gDNA) was used as positive, no-template control (NTC) with distilled water as negative control. (C) 19-day old *gfa1*_*hyp*_ *atc-2* plants in comparison to wild-type, *atc-2* and *gfa1*_*hyp*_ plants. Scale bar: 1 cm. (D) Representative images after flowering induction of 25-day old wild-type, *atc-2* and 32-day old *gfa1*_*hyp*_ *atc-2* plants at 23°C LD condition. Scale bar: 1 cm For all plant pictures brightness was equally increased in Adobe Photoshop. (E, F) Flowering time at 0.5 cm shoot elongation at 23°C LD condition (E) or SD (F) scored as rosette leaf number (see also Table S2C, H). (G) Frequency distribution of flowering wild-type, *pGFL::cGFA1/*+ and *atc-2* at 23°C under constant light (CL; black), long-day (LD; grey) and short-day (SD; green) condition (see also Table S2 D, C, H) The mean of the flowering time is indicated by an arrowhead for each light condition.

Our results thus show, that GFA1 constitutes a powerful spliceosomal switch required to implement SD-LD transition. Intriguingly, defects in BRR2a, which is an integral component of U5 snRNP interacting with GFA1 in *Arabidopsis* (Liu M. *et al*.), result in early rather than late flowering, caused by a reduction of *FLC* expression and defects in *FLC* splicing (Mahrez *et al*.). Together with the results presented in this study, a picture is emerging according to which different flowering decisions are affected by changes in spliceosome composition. These can in turn modulate the readout of a battery of flowering relevant genes in a potent and substrate specific manner.

## Supporting information

Table 1 RNASeq Data

Table 2 Flowering Assays

Table 3 Oligonucleotides

## Acknowledgements

We thank Markus Schmid for providing seeds. We thank Amelia Dolores Megía Guerrero, René Kristall, Katrischa Hennekens and Svea Küpper for technical support. C.L. and D.W. were supported by the Max Planck Society.

## Material and Methods

### Growth conditions

Transgenic plant lines were generated in *Arabidopsis thaliana* Col-0 plants. The T-DNA insertion line for *atc-2* (SALK_021669C) in Col-0 background was ordered from the Arabidopsis Biological Research Center (NASC stock number N665493). *flm-3* (NASC stock number N641971) and *cFLM-δ* OX transgenic lines were obtained from Markus Schmid laboratory (SLU Uppsala). Seeds used for RNA-seq and some RT-PCR (Fig.2D and Fig. S4A, B) were sterilized overnight in a vacuum pump using chlorine gas. Chlorine gas was produced by dropping 37% Hydrochloric acid (Sigma-Aldrich) into ∼10% Sodium hypochlorite solution (Sigma-Aldrich). For all other experiments seeds were sterilized by incubation in 80% ethanol (Carl Roth). Ethanol was exchanged 2-3 times and the total incubation time was maximum 20 minutes.

Sterile seeds were plated on half-strength Murashige-Skoog plates (0.23% MS basal salt mixture (Duchefa), 0.05% MES (Carl Roth), 1% sucrose (Carl Roth), 0.8% agar (Serva Electrophoresis), pH 5.9 with KOH), stratified at 4°C for 2 days, and grown in a growth chamber. Depending on the experiment, continuous light (LL) (24h light/0h dark; 170µmol m^-2^ s^-1^), long-day (LD) (16h light/8h dark; 170µmol m^-2^ s^-1^) or short-day (SD) (at 23°C16h light/8h dark; 170µmol m^-2^ s^-1^) conditions at 23°C or 27°C were applied. After 7 days plants were transferred to soil.

### Molecular cloning

All oligonucleotides are listed in Supplementary Table S3. The *pGFA1::cGFA1* construct corresponds with the previously published *pCLO::cCLO* construct (Moll *et al*.). To generate the *pGFL::cGFA1* construct the *pCLO* sequence was replaced with a 1.2kb genomic fragment upstream of the *GFL* coding region amplified from Col-0 DNA.

To generate the splicing reporter cassette *pAt5g40260::NLS_GFP* the promoter (Yu *et al*.) was PCR amplified from L*er* genomic DNA and placed in the pGreenIIBar vector. *NLS* amplified from *pLIS::NLS_GUS* (Groß-Hardt *et al*.) and *eGFP-*reporter lacking a start codon were inserted into pGIIBar*-pAt5g40260*. For cloning the minigenes *LIS* ^*e-i-e*^, *MEA* ^*e-i-e*^, *HD2B* ^*e-i-e*^, *SR1* ^*e-i-e*^ and *LIS* ^*e-i-e*+*1*^ were PCR amplified from L*er* DNA and inserted between *NLS* and *eGFP*. The point mutation that led to the disruption of the 5’-splice site in *LIS* ^*e*-i-e*^ was inserted by site-directed mutagenesis (Kunkel *et al*.). For plasmid and PCR product sequencing we used the LGC genomics GmbH Sanger sequencing service (LGC Ltd.). *Arabidopsis thaliana* (Col-0) was transformed by *Agrobacterium tumefaciens* assisted floral dipping (Clough *et al*.). Plants were selected using 150mg/l glufosinate solution (Bayer).

### RNA isolation, RT-PCR, qRT-PCR and Illumina sequencing

Total RNA was isolated from 30-100mg plant tissue using GeneMATRIX universal RNA Purification Kit (Roboklon). Additionally, 1 unit of RNase-Free DNaseI (Thermo Fisher Scientifc) was added to each sample. For the detection of splicing defects in *gfa1*_*hyp*_ plants (Fig. 2D and Fig. 4A-B) mRNA was isolated using Dynabeads® mRNA purification Kit for mRNA Purification from Total RNA preps (Thermo Fisher Scientific). Reverse transcription was performed with 1-5 µg total RNA using either the SuperScript® Reverse Transcription (Invitrogen) (Fig. 2D and Fig. 4A-B) or the First Strand cDNA Synthesis Kit (Thermo Fisher Scientific). For qRT-PCR we used 2µg total RNA isolated from whole rosettes without visible flower buds with AmpliTaq Gold® DNA Polymerase with Buffer II and MgCl_2_ (Thermo Fisher Scientific) according to the manufacturer’s instructions. For qRT-PCR primer and probe design we used the “Universal ProbeLibrary Assay Design Center” (Roche Holding AG) for *Arabidopsis*, to design an intron spanning assay (Table S3).

RT-PCRs were performed using either the HiFi PCR Mix (Fermentas) (Fig. 2D and Fig. 4A-B) or the lDreamTaq DNA Polymerase Kit (Thermo Fisher Scientific) according to the manufacturer’s instructions. Samples for Illumina sequencing were obtained from 12-day old Col-0 *pGFL::GFA1* (n=2) and *gfa1*_*hyp*_ seedlings (n=2). After DNase treatment, samples were prepared for sequencing using TruSeq RNA Sample Preparation Kit v2 (Illumina). Multiplex 101bp paired-end sequencing was performed by Christa Lanz (Genome Center, Max Planck Institute for Developmental Biology) using a Hiseq2000 with cBot platform (Illumina). The concentrations of isolated RNA, mRNA and cDNA were determined fluorometrically using Qubit^®^ 2.0 Fluorometer (Thermo Fisher Scientific). cDNA integrity was verified by an Agilent BioAnalyzer 2100 using DNA 1000 Chip (Agilent Technologies).

For 24h qPCR experiments, total RNA was isolated from up to 100 mg plant tissue from seedlings without visible shoot elongation using GeneMatrix universal RNA Purification Kit (Roboklon). Reverse transcription was performed with 1 µg total RNA using the First Strand cDNA Synthesis Kit (RevertAid reverse transcriptase, Thermo Fisher Scientific) and quantitative RT-PCR was performed using the Applied Biosystems StepOne Real-Time PCR system (Applied Biosystems) with TaqMan Fast Universal PCR Master Mix for TaqMan assays (#4352042 Applied Biosystems) and TaqMan Probes (Thermo Fisher Scientific, Table S3) for analysis.

### RNA-seq Data Analysis

In total, we retrieved 35M, 48M, 32M and 28M total reads for the two *pGFL::cGFA1/*+ samples and the two *gfa1*_*hyp*_ samples, respectively. Quality trimming and sequence filtering were carried out using FASTX-toolkit (v0.0.13, Hannon, G.I. (2010) FASTX-Toollkit available at http://hannonlab.cshl.edu/fastx_toolkit) with a q-value threshold of 33. Trimmed and cleaned reads were then used as input for mapping against the TAIR10 reference genome sequence of *Arabidopsis thaliana* using GSNAP (Genomic Short-read Nucleotide Alignment Program, Wu T.D. *et al*.). Duplicated reads were removed using MarkDuplicates.jar from Picard tools (v.2.18.2, available at http://broadinstitute.github.io/picard/)). Gene counts were extracted using HTSeq python tool (available at https://htseq.readthedocs.io/en/release_0.9.1/, Anders *et al*.) and TAIR10 genome annotation in gff format. Differential gene expression analysis was performed in R with edgeR, (v.3.0.8 Robinson *et al*. and McCarthy *et al*.) using whole gene counts as input and TMM (Trimmed Mean of M-values) normalization method. Genes were considered differentially expressed if their whole gene counts had an FDR (False Discovery Rate)<0.05 and log_2_FoldChange>1. Testing for DEGs was done using the negative binomial distribution model as part of the generalised linear model (GLM). For differential intron expression analysis, the same method was applied, but instead of exon counts the counts for all introns of each gene were used. RNA-seq data were visualized using Genome View N29 (Abeel *et al*.). Coverage plots consists of three areas, purple and orange show the reads mapping forward and reverse in paired-end sequencing, respectively. Green areas are mirrored along the axis and indicate the sum of all reads. The RNA sequencing data is available under ArrayExpress accession E-MTAB-14113.

### Phenotypic analysis and flowering time measurement

Flowering time point was determined as 0.5 cm shoot elongation. Rosette and cauline leaf number were determined without any tool, except for Table S2E, where we used a Leica S4E stereomicroscope. Genotype was determined in the generation before. *gfa1*_*hyp*_ plants were selected by short root, serrated and curled leaf phenotype during transfer to soil. *gfa1*_*hyp*_ genotype was randomly confirmed in a random sample of plants by PCR upon completion of the flowering assays. For root length measurements plates were imaged 7 days (Fig.S1) or 8 days (Fig.S11) after transfer to the growth chambers and the root lengths determined using ImageJ (Schneider *et al*.). For analysis of mature female gametophytes, the oldest closed flower of a given inflorescence was emasculated and harvested 2.5-3 days later. The ovaries were dissected in 5% glycerol (Carl Roth) and immediately analyzed using a Leica DMI6000b epifluorescence inverted microscope (Leica Microsystems), equipped with GFP, YFP, DAPI and DsRed filter cubes.

