## Supplemental Figures for "The spliceosomal component GAMETOPHYTIC FACTOR 1 (GFA1) regulates a key photoperiodic switch"

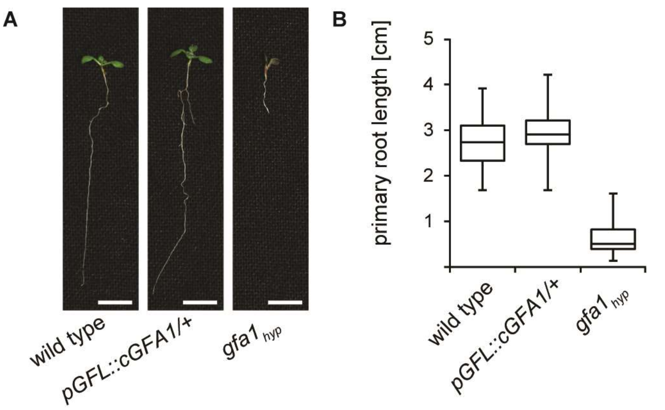


**Figure S1: Seedling phenotype and root length comparison of *gfa1*$_{hyp}$ plants.** (A) 7-day old seedlings and (B) root length measurement of wild-type (n=131), *pGFL::cGFA1/+* (n=131) and *gfa1*$_{hyp}$ plants (n=126). Scale bar: 0.5 cm. For all plant pictures brightness was increased by 120 points in Adobe Photoshop.

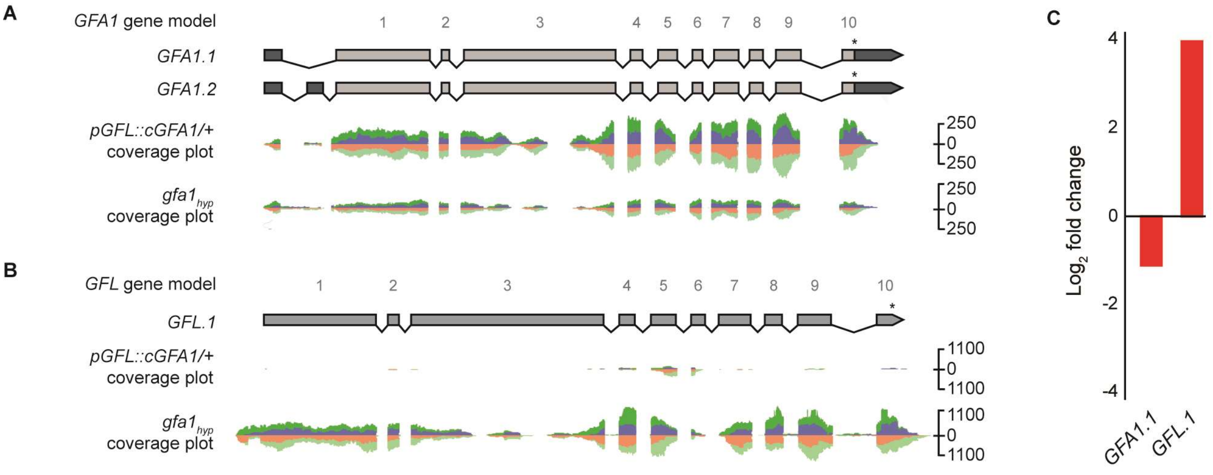


**Figure S2: *gfa1*$_{hyp}$ seedlings show differential transcript expression of *GFA1* and *GFL*.** TAIR10 gene model, RNA-seq coverage and $Log_2$ fold change in 12-day old *pGFL::cGFA1/+* and *gfa1*$_{hyp}$ seedlings for *GFA1* (A) and *GFL* (B). Introns are represented by black lines and exons by light grey boxes. UTRs represented by dark grey boxes. Stop-codon labeled by asterisks. Reads are colored by forward (dark green/orange) and reverse strand (purple/light green) for sense and antisense transcripts, respectively. (C) $Log_2$ fold change of mRNA expression level of *GFA1* and *GFL* determined by RNA-seq. Red bars displays significantly changes in mRNA expression in *gfa1*$_{hyp}$ seedlings in comparison to mRNA expression level in *pGFL::cGFA1/+* seedlings (FDR<0.05 and $log_2$ fold change>1).

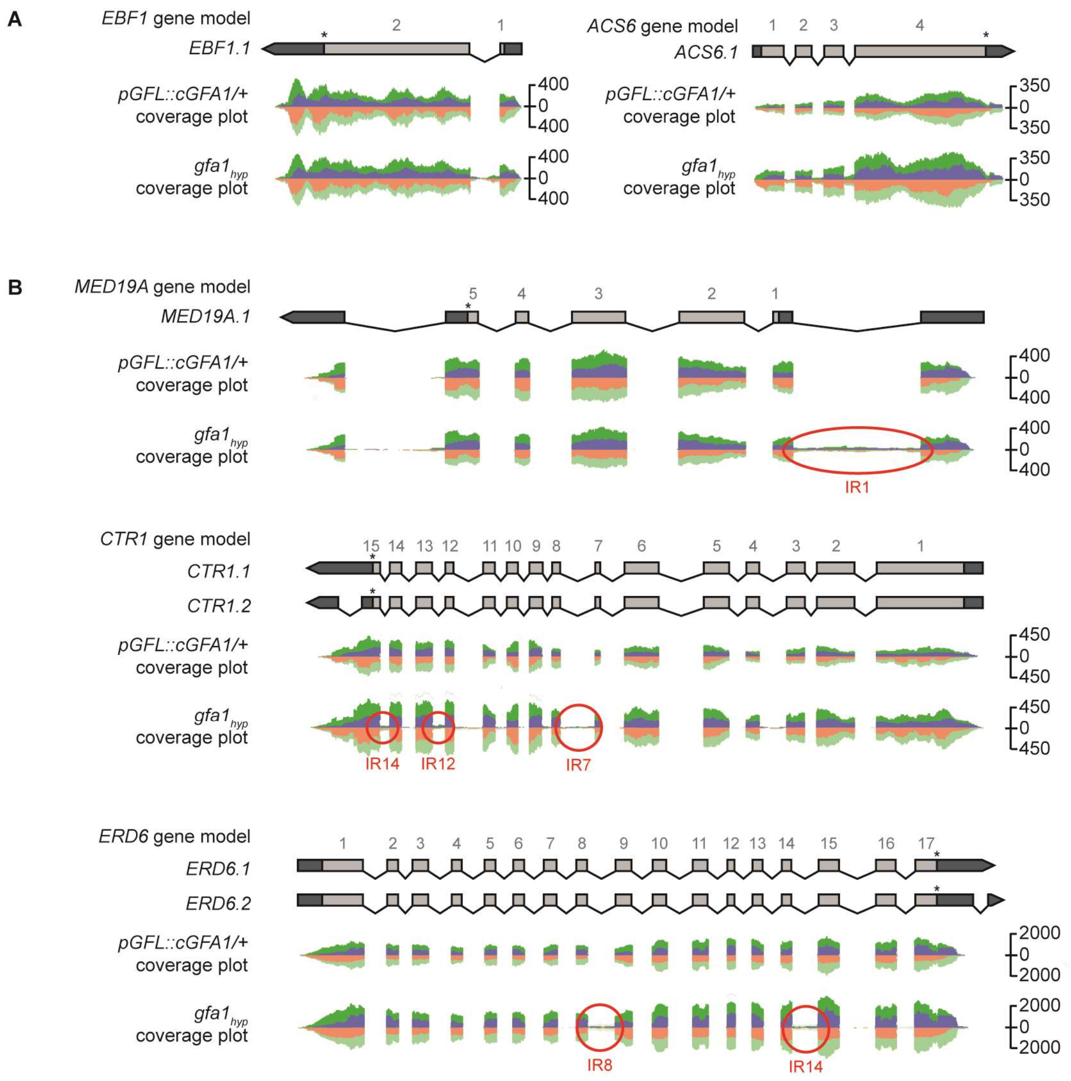


**Figure S3: *gfa1$_{hyp}$* plants exhibit substrate-specific intron retention events.** (A) TAIR10 gene model and RNA-seq coverage in 12-day old *pGFL::cGFA1/+* and *gfa1$_{hyp}$* seedlings for *EBF1* and *ACS6*. (B) TAIR10 gene model and RNA-seq coverage in 12-day old *pGFL::cGFA1/+* and *gfa1$_{hyp}$* seedlings for *MED19A*, *CTR1* and *ERD6*. Introns are represented by black lines and exons by light grey boxes. UTRs represented by dark grey boxes. Stop-codon labeled by asterisks. Reads are colored by forward (dark green/orange) and reverse strand (purple/light green) for sense and antisense transcripts, respectively. IRs are highlighted with a red circle.

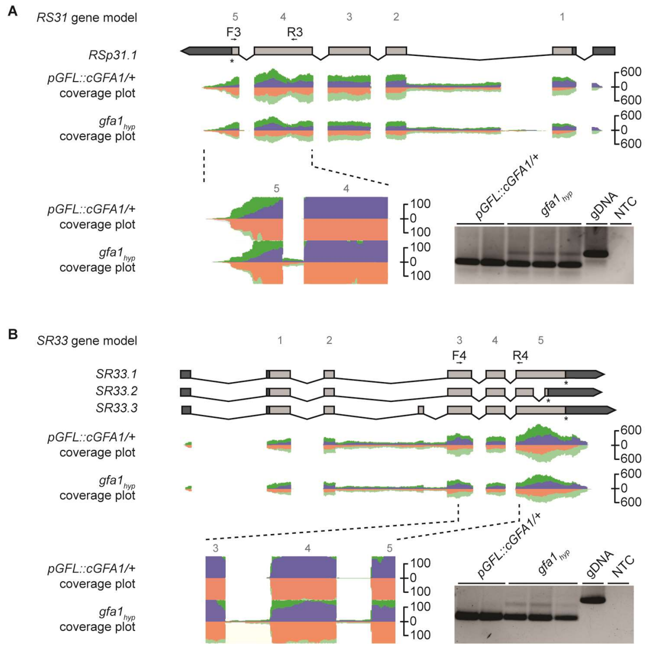


**Figure S4: Validation of intron retention events in *RS31* and *SR33* transcript in *gfa1*$_{hyp}$ plants by RT-PCR.** TAIR10 gene model and RNA-seq coverage in 12-day old *pGFL::cGFA1/+* and *gfa1*$_{hyp}$ seedlings for *RS31* (A) and *SR33* (B) . Introns are represented by black lines and exons by light grey boxes. UTRs represented by dark grey boxes. Stop-codon labeled by asterisks. Reads are colored by forward (dark green/orange) and reverse strand (purple/light green) for sense and antisense transcripts, respectively. Arrowheads indicate positions of forward (F) and reverse primers (R). Genomic DNA (gDNA) was considered as positive, no-template control (NTC) with distilled water as negative control.

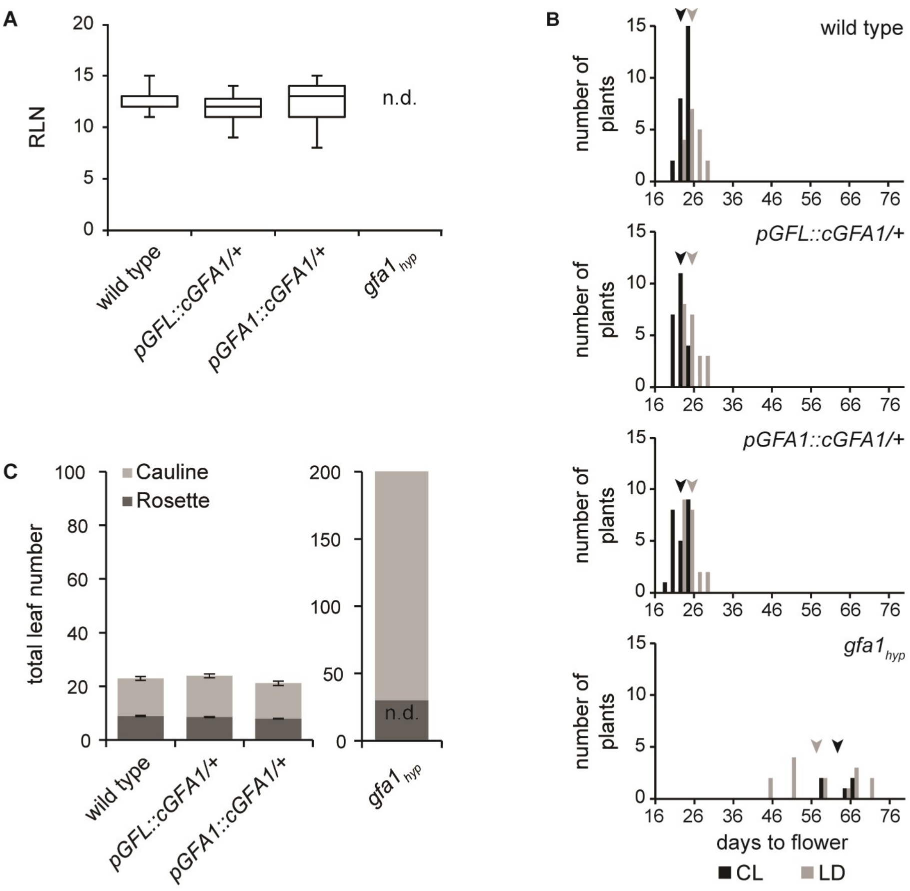


**Figure S5: *gfa1*$_{hyp}$ plants flowering phenotype at 23°C constant light (CL) and LD conditions.** (A) Flowering time (after 0.5 cm shoot elongation) at 23°C CL scored as rosette leaf number (see also Table S2D). (B) Flowering Frequency distribution at 23°C constant light (CL; black) and long-day (LD; grey) condition (see also Table S2D, A). The mean of the flowering time labeled by colored arrowheads. (C) Total leaf number at 23°C LD condition was scored as rosette leaf number and as cauline leaf number, respectively (after 0.5 cm shoot elongation) (see also Table S2E). For *gfa1*$_{hyp}$ plants total leaf number was scored 45 days after sowing and stopped when a total leaf number per plant of over 200 was reached. n.d.; indicates that clearly attribution of rosette leaves and cauline leaves was not possible.

**A**

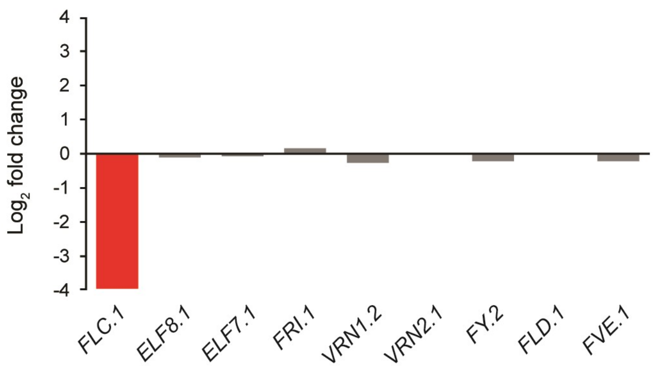


**B**

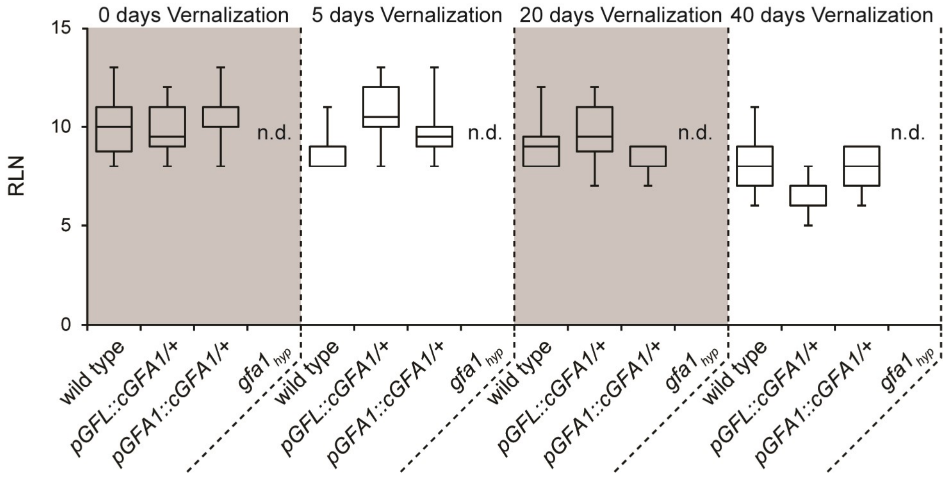


**Figure S6: Vernalization pathway analysis in *gfa1$_{hyp}$* plants.** (A) Log$_2$ fold change of mRNA expression levels of Vernalization pathway related genes determined by RNA-seq. Red bars displays significantly changes in mRNA expression in *gfa1$_{hyp}$* seedlings in comparison to mRNA expression level in *pGFL::cGFA1/+* seedlings (FDR>0.05 and log$_2$ fold change>1). (B). Effect of 0 days, 5 days, 20 days and 40 days of Vernalization on flowering time of wild type, *pGFL::cGFA1/+, pGFA1::cGFA1/+* and *gfa1$_{hyp}$* plants. Flowering time scored as rosette leaf number after 0.5 cm shoot elongation. n.d., indicates that plants did not show shoot elongation during the course of the experiment (see also Table S2F).

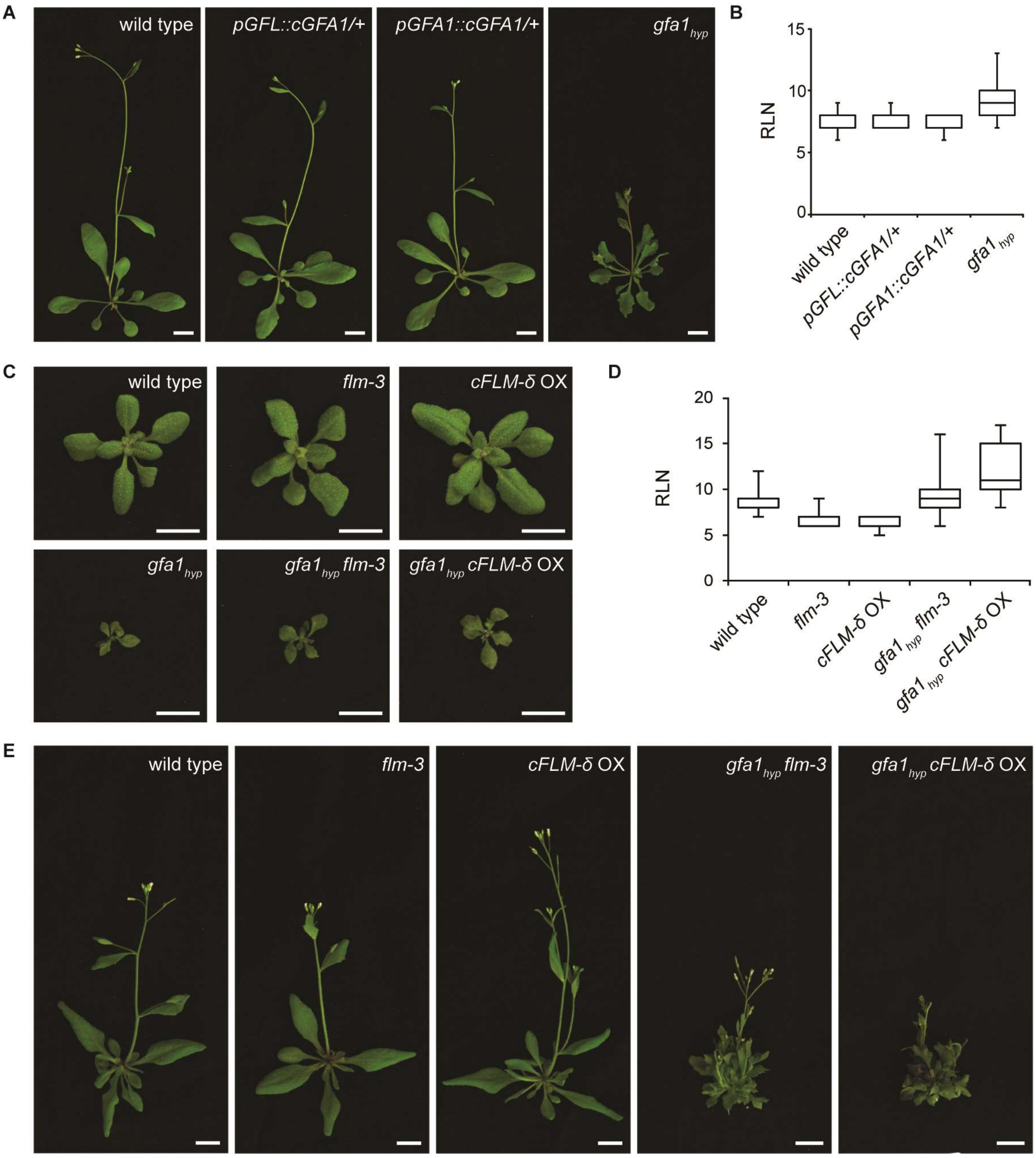


**Figure S7: Rescue of the *gfa1$_{hyp}$* flowering defect under LD condition by increased ambient temperature and by transgenic *FLM* transcript manipulation.** (A) 23-day old wild-type, *pGFL::cGFA1/+*, *pGFA1::cGFA1/+* and *gfa1$_{hyp}$* plants at 27°C LD condition after flowering induction. Scale bar: 1 cm. (B) and (D) Flowering time (after 0.5 cm shoot elongation) at 27°C LD condition scored as rosette leaf number (see also Table S2G and Table S2A). (C) 19-day old *gfa1$_{hyp}$*, *flm-3* and *gfa1$_{hyp}$ cFLM-δ* OX plants in comparison to wild-type, *pGFL::cGFA1/+*, *pGFA1::cGFA1/+* and *gfa1$_{hyp}$* plants. Scale bar: 1 cm. (E) 23-day old wild-type, *flm-3*, *cFLM-δ* OX and 40-day old *gfa1$_{hyp}$*, *gfa1$_{hyp}$ flm-3* and *gfa1$_{hyp}$ cFLM-δ* OX plants at 23°C LD condition after flowering induction. Scale bar: 0.5 cm. For all plant pictures brightness was increased by 120 points in Adobe Photoshop.

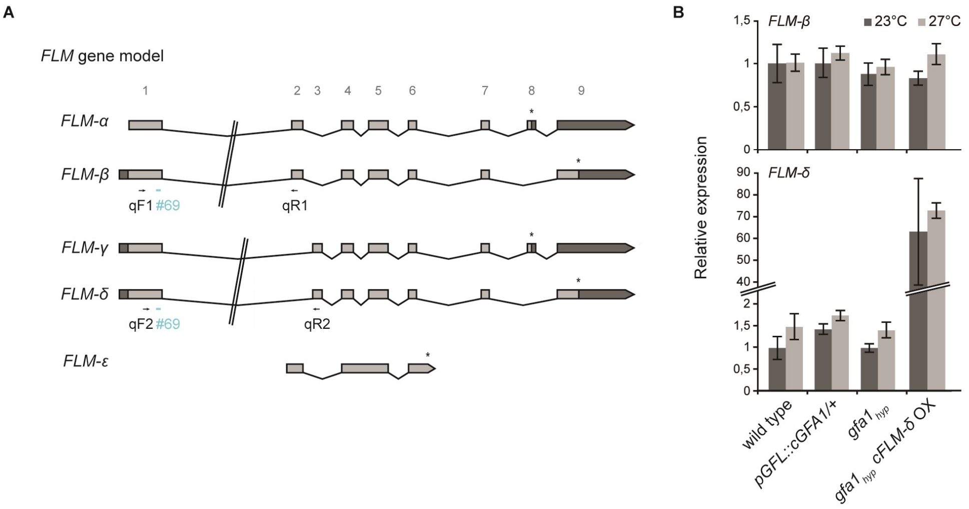


**Figure S8: Analysis of *FLM* splice variant expression in *gfa1$_{hyp}$* plants at 23°C and 27°C LD condition.** (A) TAIR10 gene model of *FLM* locus. Introns are represented by black lines and exons by light grey boxes. UTRs represented by dark grey boxes. Stop-codon labeled by asterisks. (B) qRT-PCR showing relative expression of *FLM-δ* and *FLM- β* in 20-day old wild type, *pGFL::cGFA1/+, gfa1$_{hyp}$* and *gfa1$_{hyp}$ cFLM-δ* OX plants at 23°C LD (dark grey) and 27°C LD (light grey) condition. Error bars denote s.d. of three biological replicates with three technical repetitions each. Relative expression was normalized to wild type at 23°C LD. *UBQ2* serves as house keeper. Roche probe #69 (in blue) was used to quantify both *FLM-β* (Forward primer 1, qF1; reverse primer 1, qR1) and *FLM-δ* (forward primer 2, qF2; reverse primer 2, qR2) transcript

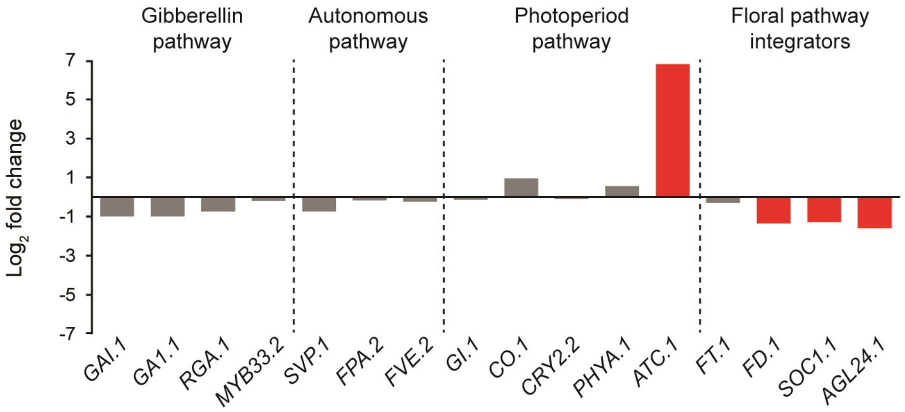


**Figure S9: Flowering pathway analysis in *gfa1$_{hyp}$* plants.** (A) $Log_2$ fold change of mRNA expression levels of genes sorted to different flowering pathways determined by RNA-seq. Red bars displays significantly changes in mRNA expression in *gfa1$_{hyp}$* seedlings in comparison to mRNA expression level in *pGFL::cGFA1/+* seedlings (FDR>0.05 and $log_2$ fold change>1).

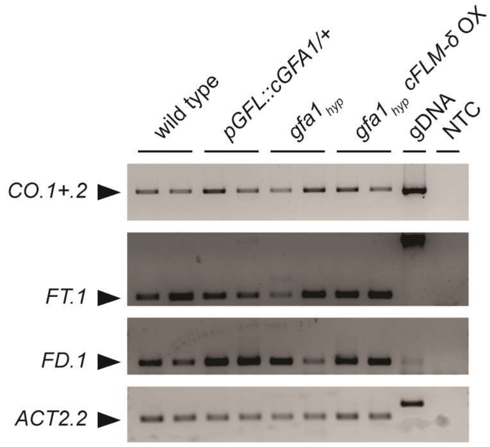


**Figure S10: Transcript analysis of floral pathway integrators.** RT-PCR on *CO.1+2, FT.1* and *FD.1* transcript expression on 20-day old wild type, *pGFL::cGFA1/+, gfa1$_{hyp}$* and *gfa1$_{hyp}$ cFLM-δ* OX plants. *ACT2.2* transcript serves as loading control. Genomic DNA (gDNA) was considered as positive, no-template control (NTC) with distilled water as negative control.

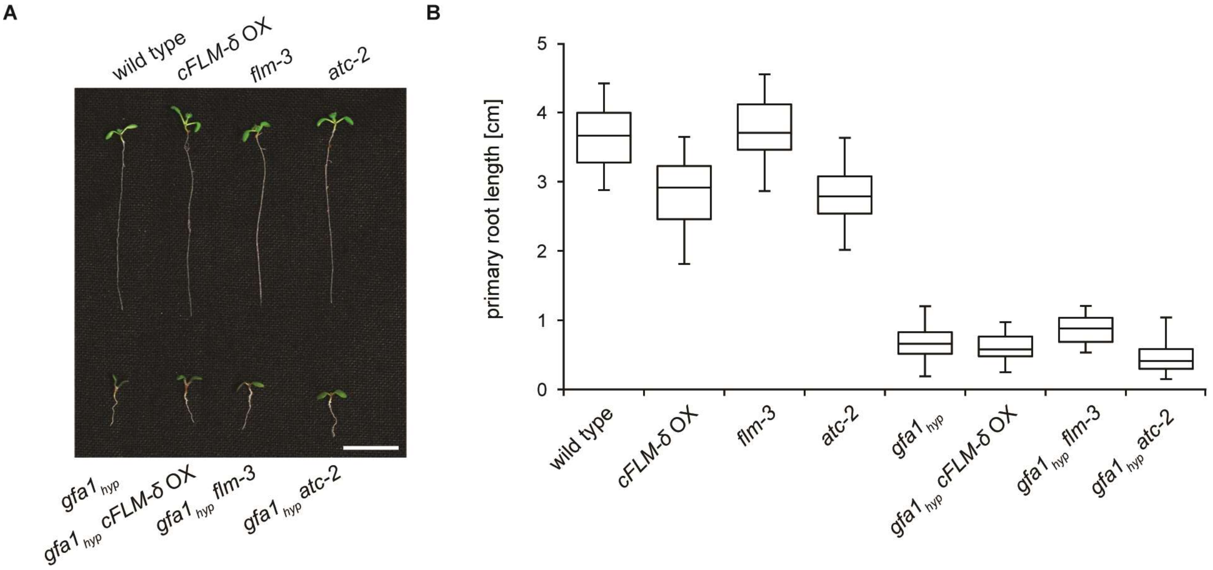


**Figure S11: Seedling phenotype and root length comparison of different *gfa1$_{hyp}$* mutant background.** (A) Representative images and (B-C) root length measurement of 8-day old wild type (n=22), *cFLM-δ* OX (n=17), *flm-3* (n=21)*, atc-2* (n=22)*, gfa1$_{hyp}$* (n=52)*, gfa1$_{hyp}$ cFLM-δ* OX (n=21)*, gfa1$_{hyp}$ flm-3* (n=20) and *gfa1hyp atc-2* (n=24) seedlings. Scale bar: 0.5 cm. For all plant pictures brightness was increased by 150 points in Adobe Photoshop.

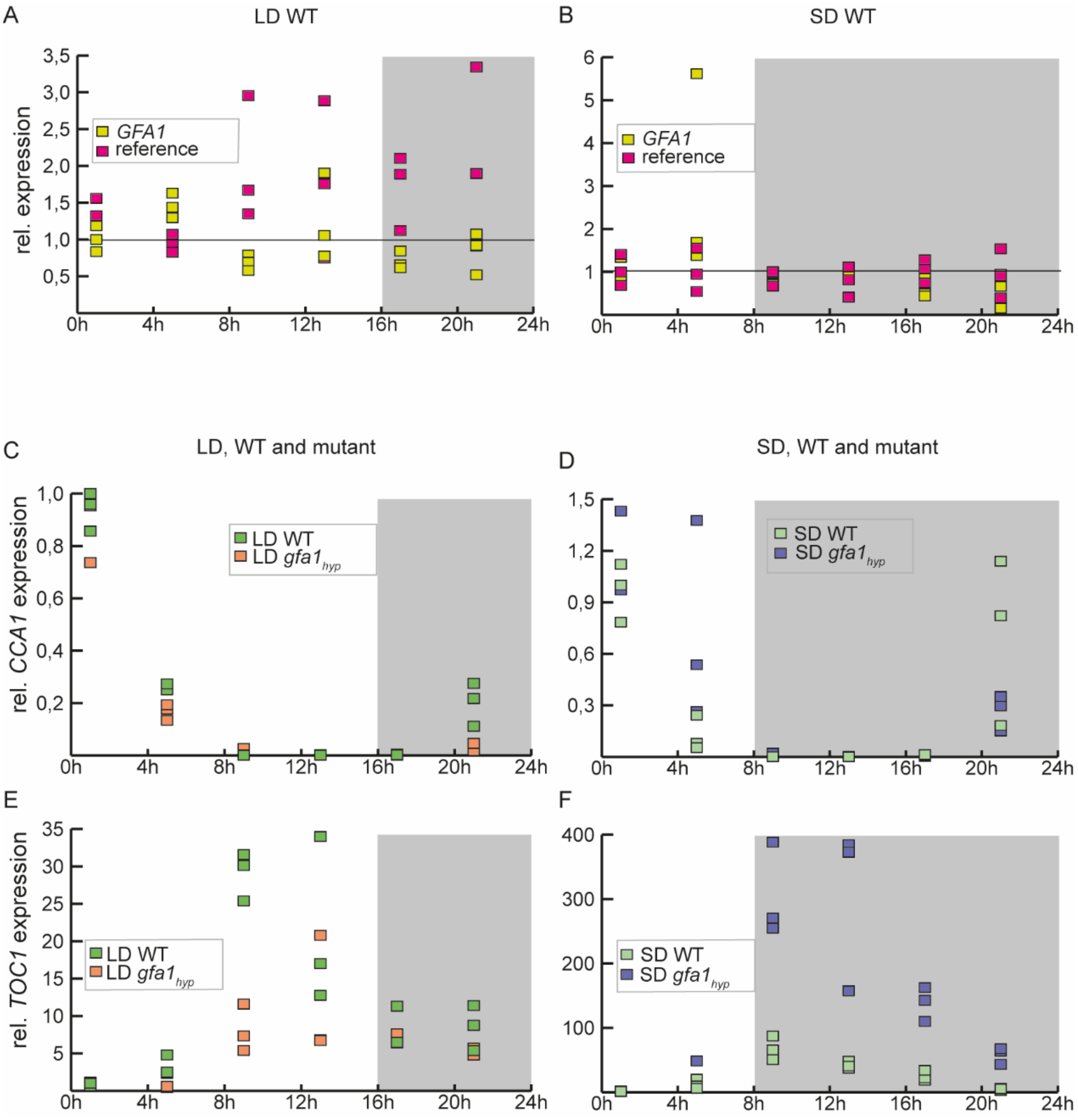


**Supplemental Figure 12: Circadian expression patterns of GFA1, CCA1 and TOC1 in WT and *gfa1*$_{hyp}$ plants** (A and B) relative expression of *GFA1* and reference (*ACT2* and *UBQ10* combined) in non-bolting wildtype plants, under SD (A) and LD (B) conditions (C and D) Relative *CCA1* expression over 24h in non-bolting wildtype and *gfa1*$_{hyp}$ plants grown under SD (C) and LD (D); (E and F) Relative *TOC1* expression over 24h in non-bolting wildtype and *gfa1*$_{hyp}$ plants grown under SD (E) and LD (F); all data as determined by qPCR, grey areas indicate dark phases, *ACT2* and *UBQ10* are used as a reference and the expression is presented relative to one data point at 1h.
